# Unravelling the molecular players at the cholinergic efferent synapse of the zebrafish lateral line

**DOI:** 10.1101/2020.07.09.196246

**Authors:** Agustín E. Carpaneto Freixas, Marcelo J. Moglie, Tais Castagnola, Lucia Salatino, Sabina Domene, Irina Marcovich, Sofia Gallino, Carolina Wedemeyer, Juan D. Goutman, Paola V. Plazas, Ana Belén Elgoyhen

**Affiliations:** Instituto de Investigaciones en Ingeniería Genética y Biología Molecular, Dr. Héctor N. Torres, Consejo Nacional de Investigaciones Científicas y Técnicas, 1428 Buenos Aires, Argentina; Instituto de Farmacología, Facultad de Medicina, Universidad de Buenos Aires, 1121 Buenos Aires, Argentina; Centro de Investigaciones Endocrinológicas “Dr. César Bergadá” (CEDIE) CONICET -FEI - División de Endocrinología, Hospital de Niños “Ricardo Gutiérrez”, 1425 Buenos Aires, Argentina

**Keywords:** lateral line, zebrafish, efferent, nicotinic receptor, calcium imaging, *Xenopus* oocytes

## Abstract

The lateral line (LL) is a sensory system that allows fish and amphibians to detect water currents. LL responsiveness to external stimuli is modulated by descending efferent neurons. LL efferent modulation aids the animal to distinguish between external and self-generated stimuli, maintaining sensitivity to relevant cues. One of the main components of the efferent system is cholinergic, the activation of which inhibits afferent activity. Since LL hair cells (HC) share structural, functional and molecular similarities with those of the cochlea, one could propose that the receptor at the LL efferent synapse is a α9α10 nicotinic cholinergic one (nAChR). However, the identity of the molecular players underlying acetylcholine (ACh)-mediated inhibition in the LL remain unknown. Surprisingly, through the analysis of single-cell expression data and *in situ* hybridization, we describe that α9, but not α10 subunits, are enriched in zebrafish HC. Moreover, the heterologous expression of zebrafish α9 subunits indicates that α9 homomeric receptors are functional and exhibit robust ACh-gated currents which are blocked by α-Bungarotoxin (α-Btx). In addition, *in vivo* Ca^2+^ imaging on mechanically-stimulated zebrafish LL HC showed that ACh elicits a decrease in evoked Ca^2+^ signals, irrespective of HC polarity. This effect was blocked by both α-Btx and apamin, indicating coupling of ACh mediated effects to SK potassium channels. Collectively, our results indicate that an α9-containing (α9*) nAChR operates at the zebrafish LL efferent synapse. Moreover, the activation of α9* nAChRs most likely leads to LL HC hyperpolarization served by the ACh-dependent activation of Ca^2+^-dependent SK potassium channels.

**Significance Statement:** Fishes and amphibians have a mechanosensory system, the lateral line (LL), which serves to detect hydromechanical variations around the animal’s body. The LL receives descending efferent innervation from the brain that modulates its responsiveness to external stimuli. LL efferent inhibition is mediated by ACh, however the identity of the molecular players at the LL efferent synapse is unknown. Here we demonstrate that a nicotinic cholinergic receptor (nAChR) composed of α9 subunits operates at the LL efferent synapse. Activation of α9-containing (α9*) nAChRs leads to LL hair cell hyperpolarization. The inhibitory signature of this process is brought about by the subsequent activation of Ca^2+^-dependent potassium SK channels, functionally coupled to α9* nAChRs.

## Introduction

The processing and encoding of external stimuli is essential for all organisms to respond appropriately to environmental cues. Fishes and amphibians have a mechanosensory system, the lateral line (LL), which senses hydrodynamic information, crucial for behaviors such as obstacle and predator avoidance, schooling, prey capture and rheotaxis (Partridge and Pitcher, 1980; Bleckmann and Zelick, 2009; McHenry et al., 2009; Suli et al., 2012; Oteiza et al., 2017).The LL comprises cell clusters, called neuromasts, composed of mechanosensitive hair cells (HC) surrounded by non-sensory cells (Metcalfe et al., 1985). Lateral line HC transmit sensory information to afferent neurons that project to the hindbrain (Metcalfe et al., 1985; Metcalfe, 1989; Liao, 2010). In addition, they are innervated by descending efferent fibers that modulate LL response to external stimuli (Metcalfe et al., 1985; Bricaud et al., 2001). During movement, LL efferent modulation aids the animal to distinguish between external and self-generated stimuli, maintaining sensitivity to relevant cues (Lunsford et al., 2019). Anatomical studies in fishes revealed two cholinergic efferent nuclei in the hindbrain, and a third dopaminergic nucleus in the forebrain (Hashimoto et al., 1970; Roberts and Russell, 1972; Zottoli and Van Horne, 1983; Tricas and Highstein, 1991; Bricaud et al., 2001). Although it is known that D1b receptors mediate neurotransmission at the dopaminergic efferent synapse (Toro et al., 2015), this information is lacking for the cholinergic one.

In mammals, the best studied efferent-HC synapse; the cholinergic medial olivocochlear (MOC) efferent system makes direct synaptic contacts with HC. The net effect of MOC activity is to hyperpolarize outer HC (OHC) (Guinan and Stankovic, 1996). This is mediated by the activation of α9α10 nicotinic cholinergic receptors (nAChRs), with very peculiar functional properties and high calcium (Ca^2+^) permeability (Elgoyhen et al., 2001; Gómez-Casati et al., 2005; Elgoyhen and Katz, 2012). Subsequent activation of Ca^2+^-dependent SK2 potassium channels drives OHC hyperpolarization (Dulon et al., 1998). Although similar molecules are probably expressed in all vertebrate efferent synapses, the mammalian α9α10 nAChR has been under positive selection, rendering a receptor with unique functional properties (Franchini and Elgoyhen, 2006; Lipovsek et al., 2012; Marcovich et al., 2020). Therefore, what has been described for the receptors at the mammalian efferent-HC synapses might not necessarily apply to other species.

Lateral line HC share structural, functional and molecular similarities with those of the cochlea. In particular, efferent stimulation to the LL and the inner ear leads to inhibition of afferent transmission (Russell Ij, 1971; Roberts and Russell, 1972; Flock and Russell, 1976; Lunsford et al., 2019; Pichler and Lagnado, 2020), thus suggesting similar synaptic mechanisms. This is most likely brought about by cholinergic efferent fibers (Dawkins et al., 2005; Zhang et al., 2018) directly contacting the base of LL HC (Dow et al., 2018), similar to what has been described for MOC efferents. Moreover, similar to cochlear OHC, LL HC have a postsynaptic cistern opposed to efferent terminals, proposed to participate in Ca^2+^ compartmentalization and/or Ca^2+^ induced Ca^2+^ release mechanisms (Lioudyno et al., 2004; Fuchs, 2014; Moglie et al., 2018; Zachary et al., 2018). These evidences suggest that the nAChR at the LL efferent synapse might be composed of α9 and α10 nAChR subunits.

To test this hypothesis we undertook a multi-pronged approach including the analysis of recent single-cell expression studies, cloning of zebrafish α9 and α10 nAChR subunits and profiling the biophysical and pharmacological properties of recombinant α9 and α9α10 receptors. In addition, we performed *in vivo* Ca^2+^ imaging of zebrafish LL HC to characterize the physiological signature of the native nAChR. We present strong evidence supporting the notion that the inhibitory signature of the LL efferent cholinergic synapse is most likely served by α9 homomeric receptors and the subsequent activation of Ca^2+^-dependent SK potassium channels.

## Materials and Methods

### Cross-study evaluation of enriched gene expression in zebrafish hair cells

Pre-processed datasets from two single-cell RNAseq (Erickson and Nicolson, 2015; Matern et al., 2018) and one microarray (Steiner et al., 2014) studies that evaluated gene expression in HC from zebrafish were used (Figure 1-1). Genes were searched by common name and accession number. For all data sets, we used the published normalized and batch-corrected gene relative expression quantification and calculated all differences as Log2 Fold Change in order to perform comparisons among studies. p-values adjusted for false discovery rate or q-values were obtained from each individual study. From the microarray dataset we used hair cells vs. mCh+, GFP+ mantle data.

### Cloning of Zebrafish nAChR cDNAs

Zebrafish RNA was isolated from 7 days postfertilization (dpf) embryos using Trizol (Thermo Fisher scientific). mRNA was reverse transcribed using polyT primers with the SuperScript™ III First-Strand Synthesis System (Thermo Fisher scientific) to obtain whole embryo cDNA. Based on the sequences reported on the Genome Reference Consortium z11, specific primers were designed to amplify whole zebrafish α9 (ENSDARG00000054680) and α10 (ENSDARG00000011113) nAChR subunits cDNAs: α9 sense (5’ - ATG AAG AGC AGT AGC AAA TAA TAA C - 3’), α9 antisense (5’ - AAT TGC AT AAG TTG TAA AC - 3’); α10 sense (5’ - ATG ATT TTA TAC TAT ATC C - 3’), α10 antisense (5’ TCA AAT GGC TTT CCC CAT TAT AAG - 3’). 35 cycles were used in both cases with an annealing temperature of 45°C and 50°C for α9 and α10 subunits, respectively. PCR products were subcloned into pCR™2.1-TOPO® TA vectors using TOPO TA Cloning Kit (Thermo Fisher scientific) and sequenced for verification of correct amplification.

### Expression of Recombinant Receptors in Xenopus laevis Oocytes

Zebrafish α9 and α10 cDNAs were sub-cloned into pSGEM vector, a modified pGEM-HE vector suitable for *Xenopus laevis* oocyte expression studies (Liman et al., 1992). All expression plasmids are readily available upon request. Capped cRNAs were transcribed *in vitro* using the RiboMAX™ Large Scale RNA Production System-T7 (Promega) from plasmid DNA templates linearized with *NheI*. Both the maintenance of *X. laevis*, and the preparation and cRNA injection of stage V and VI oocytes, has been described in detail elsewhere (Katz et al., 2000). Typically, oocytes were injected with 50 nl of RNase-free water containing 0.01–1.0 ng of cRNAs and maintained in Barth’s solution at 18°C. A 1 α9 : 2 α10 molar ratio was used to achieve expression of the heteromeric receptor.

Electrophysiological recordings were performed 2– 6 days after cRNA injection under two-electrode voltage clamp with a GeneClamp 500B Voltage and Patch Clamp amplifier (Molecular Devices). Data acquisition was performed using a Digidata 1200 and pClamp 7.0 software (Molecular Devices) at a rate of 10 points per second. Both voltage and current electrodes were filled with 3M KCl and had resistances of ∼0.5-2 MΏ. Data were analyzed using Clampfit from the pClamp 7 software suite (Molecular Devices). During electrophysiological recordings, oocytes were continuously superfused (10ml min^−1^) with normal frog saline comprised of (mM): 115 NaCl, 2.5 KCl, 1.8 CaCl_2_ and 10 HEPES buffer, pH 7.2. Drugs were applied in the perfusion solution of the oocyte chamber. V_hold_ was −70 mV except otherwise indicated.

In order to minimize the activation of the native oocyte’s Ca^2+^-sensitive chloride current (ICl_Ca_) by Ca^2+^ entering through nAChRs (Miledi and Parker, 1984; Boton et al., 1989), all experiments were carried out in oocytes pre-incubated with the membrane permeant Ca^2+^ chelator 1,2-bis (2-aminophenoxy)ethane-N,N,N’,N’-tetraacetic acid-acetoxymethyl ester (BAPTA-AM; 100 μM) for 3 h prior to electrophysiological recordings, unless otherwise stated. This treatment was previously shown to effectively chelate intracellular Ca^2+^ ions and, therefore, to impair the activation of the ICl_Ca_ (Gerzanich et al., 1994). Concentration–response curves were obtained by measuring responses to increasing concentrations of ACh.

The effects of extracellular Ca^2+^ on the ionic currents through nAChRs were studied by measuring the amplitudes of the responses to a near-EC_50_ concentration of ACh (10 μM for α9 and 300 μM for α9α10) upon varying the concentration of this cation from nominally 0 to 3 mM at a holding potential of −90 mV (Weisstaub et al., 2002). Amplitude values obtained at each Ca^2+^ concentration were normalized to that obtained in the same oocyte at 1.8 mM. Values from different oocytes were averaged and expressed as the mean ± S.E.M. These experiments were carried out in oocytes injected with 7.5 ng of an oligonucleotide (5’-GCTTTAGTAATTCCCATCCTGCCATGTTTC-3’) antisense to connexinC38 mRNA (Arellano et al., 1995; Ebihara, 1996), in order to minimize the activation of the oocyte’s nonselective inward current through a hemigap junction channel in response to the reduction of the external divalent cation concentration. Desensitization of ACh-evoked currents was evaluated via a prolonged (1 min). agonist application, at a concentration one order of magnitude above the EC_50_ for each receptor.

The percentage of current remaining 20s after the peak of the response was determined for each oocyte. Current-voltage (I-V) relationships were obtained by applying 2s voltage ramps from −120 to +50mV from a holding potential of −70mV, at the plateau response to ACh. (at a concentration one order of magnitude below the EC_50_ for each receptor).

Leakage correction was performed by digital subtraction of the I–V curve obtained by the same voltage ramp protocol prior to the application of ACh.

### Zebrafish husbandry and lines

Zebrafish (*Danio rerio*) were grown at 28.5°C on a light/dark cycle of 14:10 h, in E3 embryo medium (in mM: 130 NaCl, 0.5 KCl, 0.02 Na_2_HPO_4_, 0.04 KH_2_PO_4_, 1.3 CaCl_2_, 1.0 MgSO_4_, and 0.4 NaH_2_CO_3_). Embryos were obtained from natural spawning and bred according to guidelines outlined in The Zebrafish Book (Westerfield, 2000). For *in vivo* imaging and *in situ* hybridization experiments, 0.2 mM 1-phenyl2-thiourea (pTU) was added at 24 hours post fertilization (hpf) to prevent pigment formation. Animal experiments were done complying with the INGEBI institutional review board (Animal Care and Use Committee). Larvae were examined at 5 to 7 dpf unless otherwise stated. At these ages, sex cannot be predicted or determined, and therefore sex of the animal was not considered in our studies.

For mRNA extraction and *in situ* hybridization studies wild-type fish of the AB strain were used. For *in vivo* Ca^2+^ imaging experiments, the double transgenic line Tg [Brn3c:Gal4] [UAS:GCaMP7a] (Xiao and Baier, 2007; Muto et al., 2013) was used.

### Whole mount in situ hybridization

Embryos were fixed at 5 dpf in 4% paraformaldehyde overnight at 4°C, and stored at −20 °C in 100% methanol until use. *In situ* hybridization was performed as described previously (Thisse and Thisse, 2008). To avoid unwanted cross-reaction between nAChR genes, subunit specific probes were designed in non-conserved regions (intracellular loop of nAChR subunits) using the following primer sets: ɑ9 sense 5’-TGAAAGTGATCGAGGCCCATT-3’, ɑ9 antisense 5’-TGTTTTCCACAGACACACCCTG-3’, ɑ10 sense 5’-GGACTGCAACTGCAACATGAA-3’ and ɑ10 antisense 5’-CACCCTTCCTGTCCTCTTCCT-3’. Partial sequences of genes of interest were PCR-cloned into pCR™2.1-TOPO® using Topo TA Cloning Kit (Thermo Fisher scientific) and used as templates to perform *in vitro* transcription to synthesize sense and antisense digoxigenin (DIG)-labeled probes. Sense probes were used as negative controls. Larvae were imaged on a Nikon Eclipse E200 microscope using a Nikon E Plan 10x/0.25 objective lens. Images were acquired via a Micrometrics® 891CU CCD 8.0 Megapixel camera using Micrometrics® SE Premium imaging software.

### Sample preparation and stimulation for functional imaging

Individual Tg [Brn3c:Gal4;UAS:GcAMP7a] larvae at 5-7 dpf were first anesthetized with tricaine (0.03% ethyl 3-aminobenzoate methanesulfonate salt) and then pinned (through the head and tail) onto a Sylgard-filled recording chamber. To suppress movement, α-Btx (125 μM) was injected directly into the heart. Larvae were then rinsed with extracellular imaging solution (in mM: 140 NaCl, 2 KCl, 2 CaCl_2_, 1 MgCl_2_, and 10 HEPES, pH 7.3, OSM 310±10) without tricaine and allowed to recover. Viability was monitored by visually monitoring heart rate and blood flow.

Stimulation of neuromast HC was accomplished using a custom-made fluid jet. Pressure was applied using a 15 ml syringe and controlled through a TTL valve system (VC-6 valve controller, Warner instruments) triggered via the recording system. The output was attached to a glass pipette (inner tip diameter ~30–50 μm) filled with extracellular imaging solution and positioned parallel to the anterior–posterior axis of the fish in order to mechanically stimulate the apical bundles of HC along that axis (deflections were sustained for the duration of the stimulus, without flickering and kinocilial deflections were confirmed visually). We used the fluid jet to stimulate the HC of the two polarities by applying either negative or positive pressure. Air volume injected through the syringe was constant (5 ml) and pressure was controlled with a manometer. Two second square stimuli were delivered in order to activate HC of all sensitivities (Zhang et al., 2018; Pichler and Lagnado, 2019).

### Functional imaging

Fish were placed into a chamber on the stage of an upright microscope (Olympus BX51WI), illuminated with a blue (488 nm) LED system (Tolket) and images were acquired using an Andor iXon 885 camera controlled through a Till Photonics interface system. The focal plane was located close to the basal region of the neuromast in order to visualize the basal pole of HC. The signal-to-noise ratio was improved with a chip binning of 4×4, giving a resolution of 0.533 μm per pixel using a 60X water immersion objective. Acquisition rate was set to 6.6 frames/sec. Isradipine, α-Btx and apamin were applied in the bath and fish were pre-incubated prior to image acquisition. ACh was locally perfused throughout the image acquisition protocol. Fluorescence images were processed in FIJI (Schindelin et al., 2012; Rueden et al., 2017) and analyzed with custom-written routines in IgorPro 6.37 (Wavemetrics). Images were motion corrected using the StackReg plugin (Thévenaz et al., 1998).

Regions of interest (ROIs) were hand-drawn for each visible HC in the neuromast. The mean ΔF/F 0 (%) was calculated in every ROI for each time frame and corrected for photobleaching by fitting a line between the pre-stimulus baseline and final fluorescence. Peak fluorescence signals were detected during the mechanical stimulation of the neuromast. Further analysis was performed if the peak signal was at least 2.5 standard deviations higher than the baseline.

### Statistical analysis

For the biophysical characterization of recombinant receptors in *Xenopus laevis* oocytes all plotting and statistical tests were conducted using Prism 6 software (GraphPad Software Inc.). Concentration-response curves were normalized to the maximal agonist response in each oocyte. The mean and S.E.M. values of the responses are represented. Agonist concentration-response curves were iteratively fitted with the equation I / Imax=[A]^n^ / ([A]^n^+EC_50_ ^n^), where I is the peak inward current evoked by the agonist at concentration [A]; Imax is the current evoked by the concentration of agonist eliciting a maximal response; EC_50_ is the concentration of agonist inducing half-maximal current response and n is the Hill coefficient.

One-way repeated measures ANOVA, with a Geisser-Greenhouse correction to account for non-sphericity, was run to determine if there were statistical significant differences in responses to extracellular Ca^2+^concentrations. A Bonferroni multiple comparison test was performed to evaluate differences between group means.

For *in vivo* Ca^2+^ imaging, all experiments were performed on a minimum of 8 animals (1 neuromast per animal) and on three independent days. Plotting and statistical analysis were performed using a custom written code in Python language (Python 3.7), using pandas, Scipy, numpy, IPython, matplotlib and seaborn packages (Hunter, 2007; Pérez and Granger, 2007; McKinney, 2010; Walt et al., 2011; Virtanen et al., 2020). Normality was tested using a Shapiro-Wilk normality test. As data was not normally distributed, statistical significance between two conditions was determined by Wilcoxon matched-pair ranks test. For this kind of analysis, effect sizes were calculated as Matched Pairs Rank Biserial Correlation (MPRBC), which equals the simple difference between the proportion of favorable and unfavorable evidence (Kerby, 2014). Statistical significance is reported at α = 0.05

### Drugs

All drugs were obtained from Sigma-Aldrich, except α-Btx and apamin that were purchased from Alomone. For *in vivo* imaging experiments, drugs were brought to their final concentration in normal extracellular imaging solution with 0.1% DMSO. Isradipine, apamin and α-Btx were applied in the bath during pre-incubations and locally perfused. ACh was locally perfused.

## Results

### Cross-study evaluation of enriched gene expression in zebrafish hair cells

In order to decipher the molecular players at the cholinergic efferent LL synapse, we first studied the expression of genes that encode key molecules of efferent synapses across vertebrates: chrna9 (gene encoding the α9 nAChR subunit), chrna10 (α10 subunit) and kcnn2 (small conductance Ca^2+^-activated potassium channel 2, SK2), in LL HC. As there is evidence for the expression of other SK channels in zebrafish sensory organs (Cabo et al., 2013), we also evaluated the expression of kcnn1 and kcnn3 (genes encoding SK1 and SK3 channels, respectively). The Genome Reference Consortium Zebrafish Build 11 (GRCz11) indicates two ohnologues for kcnn1 (kcnn1a and kcnn1b) and only one copy for chrna9, chrna10, kcnn2 and kcnn3. A previous genome assembly version, GRCz10, had described two ohnologues for both chrna9 and chrna10 genes, but one of the copies of each gene (ENSDARG00000011029 and ENSDARG00000044353) have been deprecated in GRCz11.

We collected data from recently published single-cell RNA-seq and microarray studies in zebrafish HC (Steiner et al., 2014; Erickson and Nicolson, 2015; Matern et al., 2018) (Figure 1-1) and assessed the enrichment of chrna9, chrna10, kcnn1a, kcnn1b, kcnn2 and kcnn3 genes. Pre-processed data from each study (normalized, batch-corrected and with their adjusted p-values) was analyzed. The relative change in expression (Log_2_ fold change) in HC was normalized to the control sample used in each study (Figure 1-2).

Our analysis revealed that only chrna9, kcnn1a and kcnn2 transcripts are significantly enriched in HC (Figure 1). These observations were true for all data sets analyzed in the case of chrna9 transcripts and in two of the three studies evaluated, in the case of kcnn1a and kcnn2. Surprisingly, chrna10 transcripts showed no enriched expression in HC (Figure 1).

**Figure 1.**
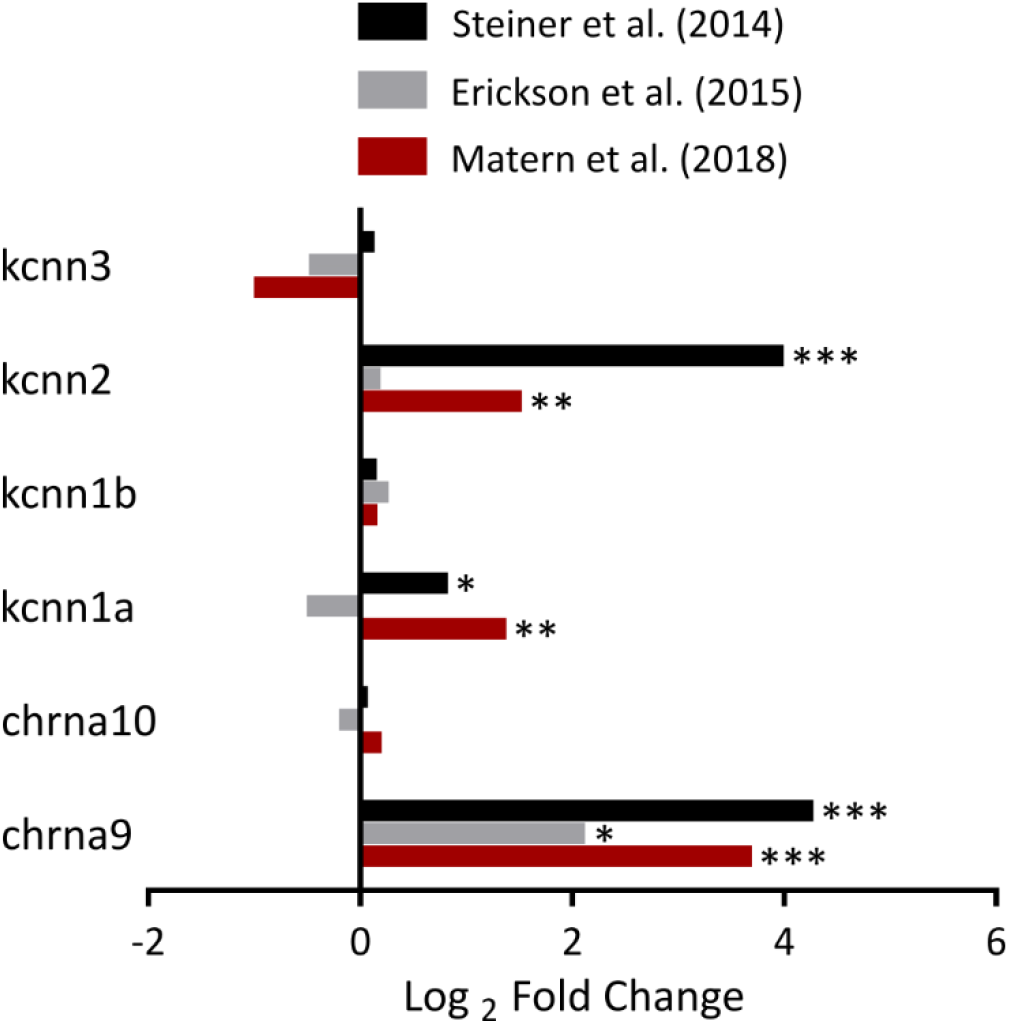
ɑ9 (but not ɑ10), SK1a and SK2 are enriched in zebrafish HC. Cross-study relative expression levels (Log_2_FC) for genes of interest. In all cases, ɑ9 (chrna9) is enriched compared to control in HC. Data from Matern et al. (2018) and Erickson et al. (2015) also show a significant enriched expression for SK2 (kcnn2) and SK1a (kcnn1a). Adjusted p-values are taken from each study (Figure1-2, *p<0.05, **p<0.005, ***p<0.00005).

To further analyze the spatial expression pattern of α9 and α10 nAChRs subunits we performed whole mount *in situ* hybridization in 5 dpf larvae. To avoid unwanted cross-reaction between nAChR genes, subunit specific probes were designed in the non-conserved intracellular loop of nAChR subunits. α9 subunit expression was localized to LL neuromasts and the otic vesicle (Figure 2), confirming its specific expression in HC. No signal was detected for the α10 subunit mRNA (data not shown).

**Figure 2.**
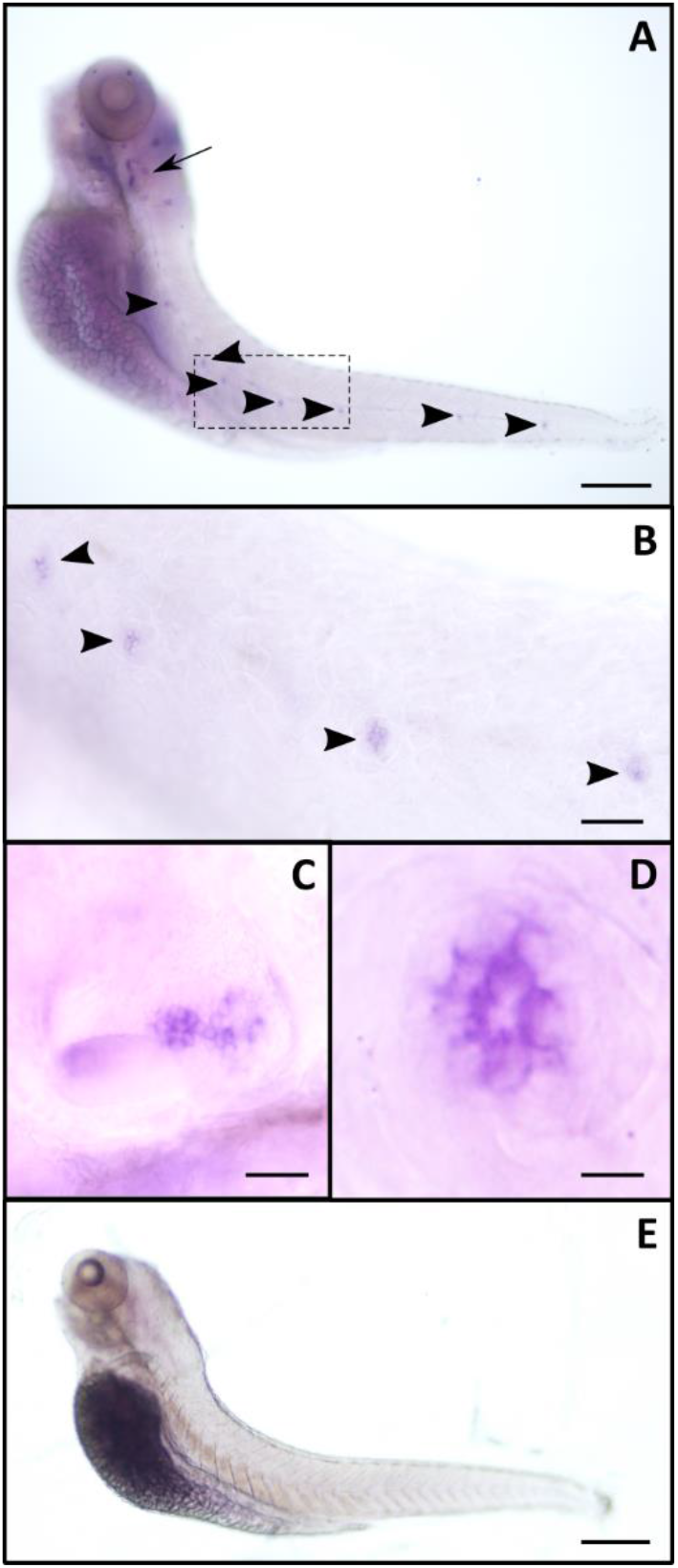
Expression pattern of zebrafish ɑ9 nAChR subunit mRNA in 5 dpf zebrafish larvae. Whole-mount *in situ* hybridization with antisense (**A, B, C** and **D**) and sense (**E**) ɑ9 riboprobes. Representative lateral views with anterior to the left and dorsal to the top, are shown. Arrow indicates the otic vesicle and arrowheads point to selected neuromasts. Large scale view of the otic vesicle (**C**) and neuromasts (**B** and **D**). dpf, days post-fertilization. Scale bars: 100 μm in **A** and **E**, 40 μm in **B**, 25 μm in **C**, 10 μm in **D**.

### Biophysical and pharmacological characterization of zebrafish recombinant α9 and α9α10 nAChRs expressed in Xenopus laevis oocytes

To determine the possible combinatorial nAChR subunit assemblies leading to functional receptors and to analyze their pharmacological and biophysical properties, we performed RT-PCR with specific primers designed to isolate full-length zebrafish α9 and α10 nAChR subunit cDNAs, and subcloned them into pSGEM vector (a pGEM-HE vector optimized for *X. laevis* oocyte expression studies (Liman et al., 1992)). *In vitro* transcribed cRNAs were injected in *X. laevis* oocytes and responses to ACh were recorded under two-electrode voltage clamp.

Previous work reported that *Rattus norvegicus* (rat), *Xenopus tropicalis* (frog) and *Gallus gallus* (chicken) α9 subunits can form functional homomeric nAChRs. In contrast, only chicken and frog α10, but not rat subunits, assemble into functional homomeric receptors (Elgoyhen et al., 1994, 2001; Lipovsek et al., 2012, 2014; Marcovich et al., 2020). Whereas zebrafish α9 subunits assembled into functional homomeric receptors leading to robust ACh-evoked currents (Imax 425.52 ± 55.00 nA, n= 28), α10 subunits could not form functional receptors under our experimental conditions (Figure 3A). Oocytes injected with both α9 and α10 zebrafish cRNAs in an equimolar proportion responded to ACh in a concentration-dependent manner with a two component ACh dose-response curve that corresponds to both homomeric α9 and heteromeric α9α10 nAChRs (Figure 3B). To favor the assembly of α9α10 heteromeric receptors and study its properties, we injected both cRNAs in a 1:2 α9:α10 ratio. ACh concentration-response curves for homomeric α9, α9α10 (1:1) and α9α10 (1:2) nAChRs are shown in Figure 3B. α9 nAChRs exhibited an EC_50_ of 11.71 μM (n=10; 95% CI 9.48 to 14.46), while the EC_50_ for α9α10 (1:2) receptors was 437 μM (n=7; 95% CI 357.2 to 534.6). It is interesting to note that, in contrast to that reported for rat receptors, α10 did not boost responses of the heteromeric α9α10 (Imax 292.05 ± 52.13 nA, n= 30), compared to the homomeric α9.

**Figure 3.**
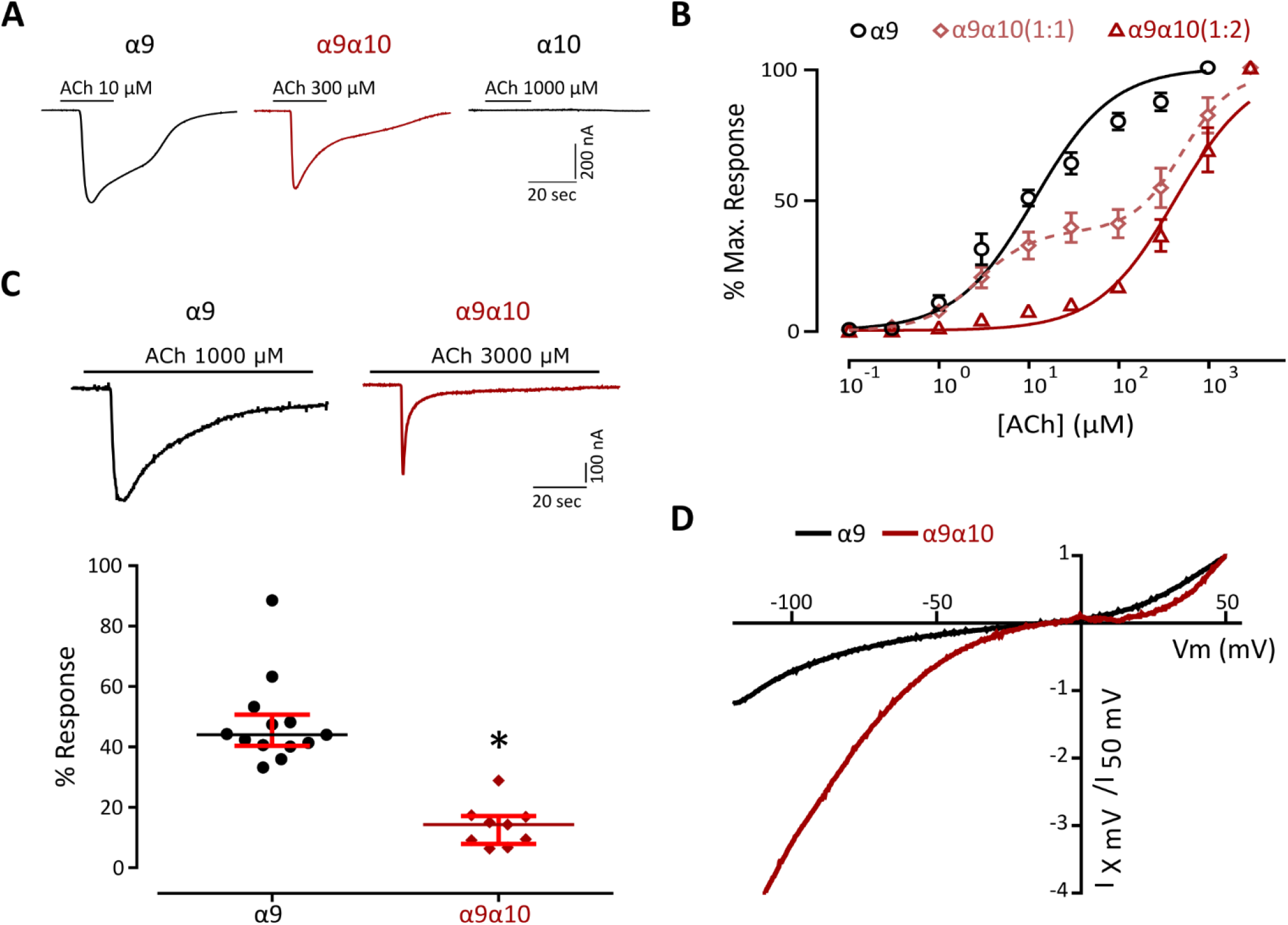
Biophysical characterization of zebrafish recombinant α9 and α9α10 nAChRs. **A**, Representative responses evoked by ACh in oocytes expressing either zebrafish ɑ9, ɑ10 or ɑ9ɑ10 (1:2) nAChRs. **B**, Concentration-response curves for zebrafish ɑ9, ɑ9ɑ10 (1:1) and ɑ9ɑ10 (1:2) nAChRs. Values are the mean ± S.E.M. Lines are best fit to the Hill equation. **C**, *Top*, Representative responses of zebrafish α9 and α9α10 (1:2) nAChRs to a 60 s application of ACh (1 order of magnitude higher than their corresponding EC_50_); *Bottom*, percentage of current remaining 20s after the peak response relative to the maximum current amplitude elicited by ACh. Lines indicate the median and IQR. Symbols represent individual oocytes (n= 13 and 9, respectively). **D**, Representative I-V curves obtained by the application of voltage ramps (−120 to +50 mV, 2s) at the plateau response to 10 μM ACh for both zebrafish α9 and α9α10 (1:2) nAChRs. Values were normalized to the agonist response at +50 mV for each receptor.

### Desensitization profile

A key feature of nAChRs is their desensitization after prolonged exposure to ACh (Quick and Lester, 2002). Zebrafish α9 and α9α10 receptors exhibited different desensitization profiles (Figure 3C). While in the case of the α9 nAChR a median of 44% (IQR: 40.36-50.75 %) of remaining current was observed 20 seconds after the peak response to 1 mM ACh, a median of 14.29% (IQR: 7.91-17.15 %) was observed for α9α10 receptors 20 seconds after the peak response to 3 mM ACh, indicating a faster desensitization (*p= 7.09e-06, U = 0.0, Mann-Whitney test).

### Current-voltage relationship

Another distinctive feature that varies among α9α10 nAChRs is their current-voltage (I-V) relationship. Rat α9α10 receptors show a significant outward current at depolarized potentials and a greater inward current at hyperpolarized potentials (Elgoyhen et al., 2001). Chicken α9α10 nAChRs exhibit outward currents similar to their rat counterparts but smaller inward currents (Marcovich et al., 2020), and frog α9α10 receptors show an I-V profile with strong inward rectification and almost no outward current at depolarized potentials (Marcovich et al., 2020). Zebrafish α9α10 nAChRs also showed an unique I-V profile (Figure 3D), exhibiting considerable outward currents at depolarized potentials, similar to chicken α9α10 and rat α9 and α9α10 receptors (Elgoyhen et al., 2001). At hyperpolarized potentials, although with different amplitudes, both receptors exhibited inward rectification similar to their frog counterpart (Marcovich et al., 2020).

### Ca^2+^ contribution to ACh-evoked responses

Ca^2+^ entry through α9α10 nAChRs is key for the function of the MOC-HC synapse, since the subsequent activation of Ca^2+^-dependent potassium channels ultimately leads to the hyperpolarization of the HC. Ca^2+^ permeability of α9α10 nAChRs is not uniform across species (Lipovsek et al., 2012, 2014; Marcovich et al., 2020). In order to assess Ca^2+^ flux through zebrafish α9 and α9α10 nAChRs we analyzed the contribution of the *Xenopus* oocytes endogenous *ICl*_*Ca*_ (Miledi and Parker, 1984; Boton et al., 1989) to ACh-evoked responses. In oocytes expressing a recombinant receptor with high Ca^2+^ permeability, the *ICl_Ca_* is strongly activated upon ACh application (Barish, 1983). Incubation of oocytes with the membrane-permeant fast Ca^2+^ chelator BAPTA-AM subsequently abolishes the Cl^−^component of the total measured current (Gerzanich et al., 1994). Responses to ACh showed a strong reduction after BAPTA incubation (Figure 4A, α9: 45.05 ± 5.87 % of current remaining, n = 15; α9α10: 38,97 ± 5.48 % of current remaining, n= 12), indicating a significant Ca^2+^ contribution to ACh-evoked responses for both receptors.

**Figure 4.**
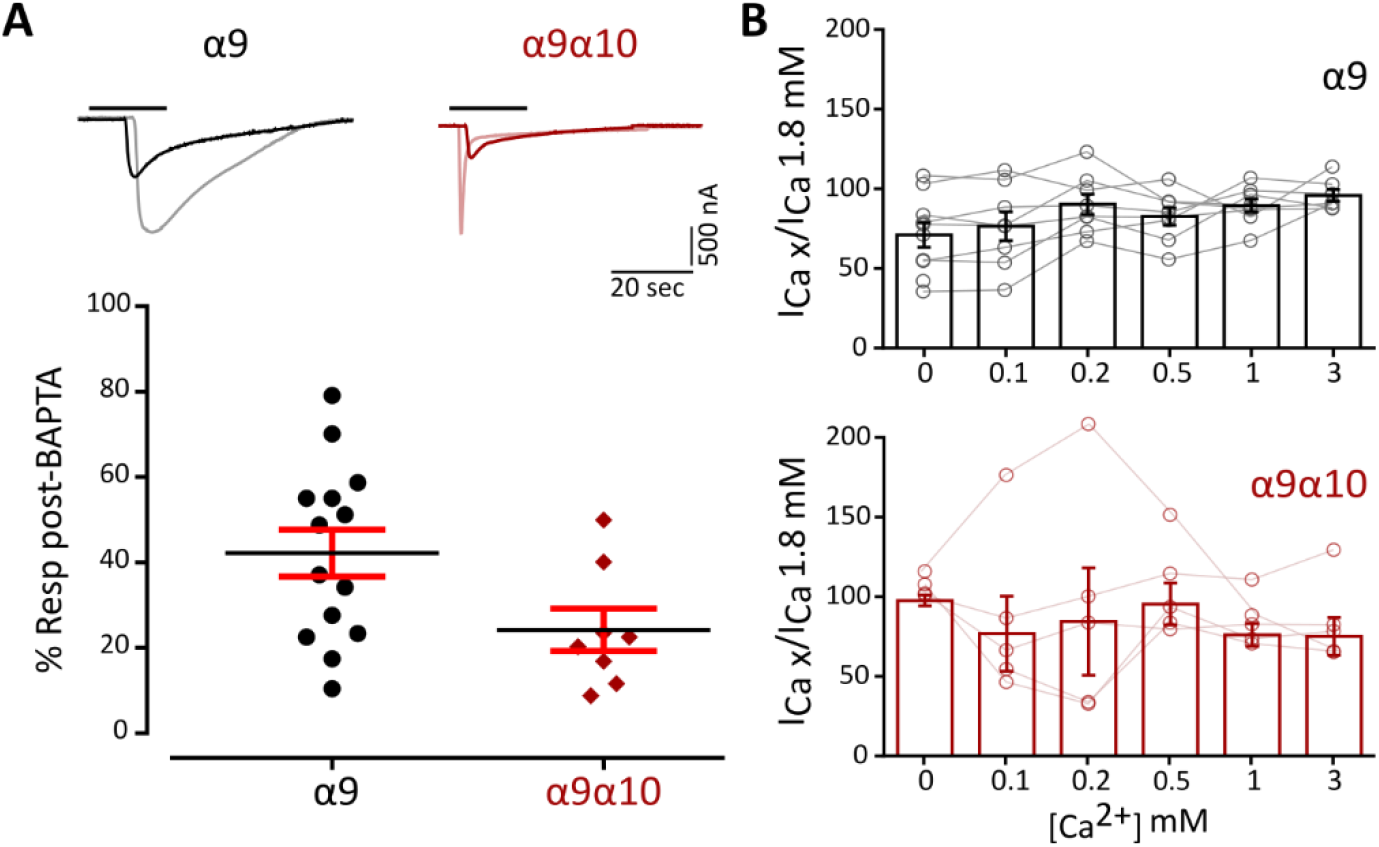
Zebrafish α9 and α9α10 nAChRs have a high Ca^2+^ contribution to the total inward current and are not modulated by extracellular Ca^2+^ **A**, *Top*, Representative responses to a near-EC_50_ concentration of ACh (10 μM for α9 and 300 μM for α9α10) in oocytes expressing zebrafish α9 and α9α10 nAChRs before (light colors) and after (solid colors) a 3-h incubation with BAPTA-AM. *Bottom*, Percentage of the initial response remaining after BAPTA incubation. Lines indicate the median and IQR. Symbols represent individual oocytes (n= 14 and 8, respectively), **B**, ACh response amplitude as a function of extracellular Ca^2+^ concentration (*Top*, α9; *Bottom*, α9α10). ACh was applied at a near EC_50_ concentration (10 μM for α9 and 300 μM for α9α10). Current amplitudes recorded at different Ca^2+^ concentrations in each oocyte were normalized to the response obtained at 1.8 mM Ca^2+^ in the same oocyte. Vhold: −90 mV. Bars represent mean ± S.E.M, (n= 8 for α9 and 5 for α9α10).

### Modulation of ACh-evoked responses by extracellular Ca^2+^

α9α10 nAChRs from rat, chicken, and frog exhibit differential modulation by extracellular Ca^2+^ (Weisstaub et al., 2002; Marcovich et al., 2020). Rat α9α10 receptors are both potentiated and blocked by extracellular Ca^2+^, whereas in the case of frog and chicken α9α10 nAChRs ACh responses are only potentiated by this cation. Moreover, rat α9 receptors are only blocked by extracellular Ca^2+^. To evaluate Ca^2+^ modulation of zebrafish α9 and α9α10 receptors, responses to near EC_50_ concentrations of ACh were recorded in normal Ringer’s solution at different extracellular Ca^2+^ concentrations and normalized to the response at 1.8 mM Ca^2+^. Strikingly, neither the homomeric α9 nor the heteromeric α9α10 nAChR responses to ACh were modulated by extracellular Ca^2+^ (Figure 4B).

### Pharmacological characterization

A hallmark of α9 and α9α10 nAChRs is their peculiar pharmacological profile, which includes α-Btx and strychnine as potent inhibitors. We studied the effect of both drugs on α9 and α9α10 nAChRs expressed in *Xenopus* oocytes. As shown in Figure 5, pre-exposure of oocytes for 1 min with 100 nM α-Btx before the co-application of 10 μM (α9) or 300 μM (α9α10) ACh reduced agonist-evoked response by 93.70 ± 2.17% (n = 2) and 85.16 ± 6.86% (n= 2), respectively. In both cases the effect of α-Btx was completely reversed by washing the oocytes with frog saline solution for 5 min. Similar block of ACh responses was obtained with strychnine (data not shown).

**Figure 5.**
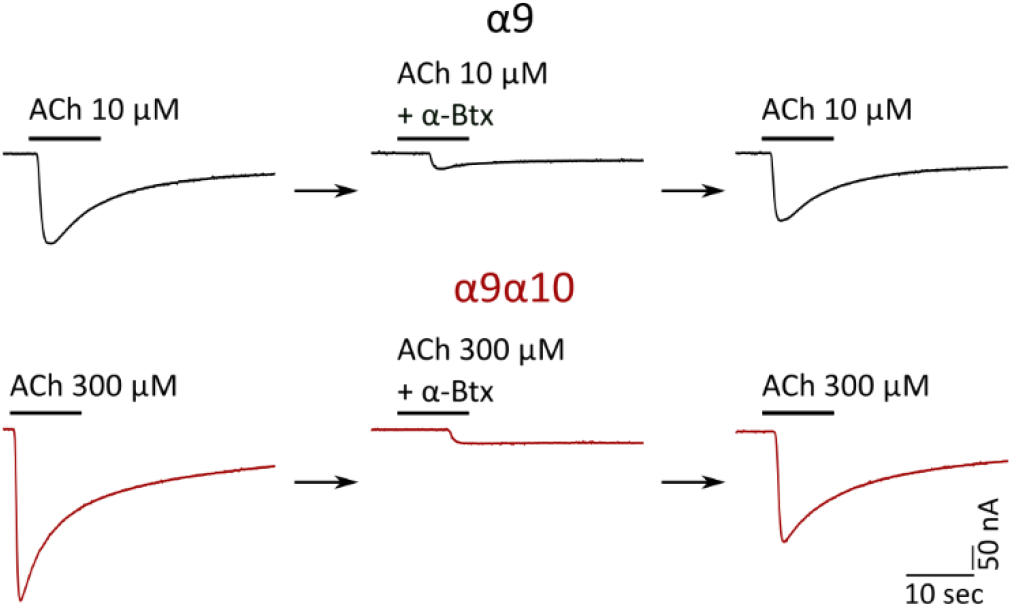
Zebrafish α9 and α9α10 nAChRs are reversibly blocked by ɑ-Btx. Responses to 10 μM (α9) or 300 μM (α9α10) ACh either alone, in the presence of 100 nM α-Btx or after washing with control bath solution for 5 min, in oocytes expressing zebrafish α9 or α9α10 nAChRs are shown. Oocytes were pre-incubated with 100 nM α-Btx for 1 min prior to the addition of the agonist. α-Btx inhibited ACh responses by 93.70 ± 2.17% (n = 2) and 85.16 ± 6.86% (n= 2), respectively.

### In vivo functional Ca^2+^ imaging

To characterize the physiological signature of the native nAChR present at the zebrafish LL efferent synapse, we performed *in vivo* Ca^2+^ imaging in transgenic Tg [Brn3c:Gal4;UAS:GcAMP7a] zebrafish larvae that specifically express the genetically encoded Ca^2+^ sensor GcAMP7a in HC. We mechanically stimulated LL HC with saturating stimuli in both rostral and caudal orientations, by applying positive and negative pressure through a pulled glass pipette respectively, to elicit the activation of the mechanotransduction channel and thus produce depolarization of HC and subsequent opening of voltage-gated Ca^2+^ channels (Cav 1.3a) (Zhang et al., 2018; Pichler and Lagnado, 2019). We measured peak fluorescence signals derived from mechanically evoked Ca^2+^ influx through Cav1.3a channels as a proxy of the electrical state of HC (Figure 6A and B). Posterior LL HC are variable in their function and signal transduction properties (Zhang et al., 2018; Pichler and Lagnado, 2019). We therefore chose as our experimental unit single HC that were selective for one polarity; that is HC that showed robust activation either by positive or negative deflections (e.g cells 1,2,3,4,5,8 and 9 in Figure 6B). HC exhibited robust and stable Ca^2+^ signals over two trials with the same stimulation after 1 minute (Figure 6C, 1° stim: Mdn ΔF/F0= 0.858 IQR: 0.472-1.504 Vs. 2° stim: Mdn ΔF/F0= 0.876 IQR: 0.475-1.503, n= 113, W= −835, p= 0.2317, matched pairs rank biserial correlation (MPRBC)= 0.129 Wilcoxon matched-pairs signed rank test). Consistent with previous findings (Sheets et al., 2012; Zhang et al., 2018), HC pretreated with the Cav1.3 antagonist isradipine (10 *μ*M) showed a significant decrease in Ca^2+^ entry levels with respect to control conditions (Figure 6D, Ctrl: Mdn ΔF/F0= 0.743 IQR: 0.249-1.086 Vs. Isr: Mdn ΔF/F0= 0.209 IQR: 0.079-0.433, n = 23, W= −258, p = 8.726e-05, MPRBC= 0.935; Wilcoxon matched-pairs signed rank test). This finding reveals that under our experimental conditions a large proportion of the total change in fluorescence intensity after mechanical stimulation can be attributed to Ca^2+^ influx through Cav1.3a channels. The remaining change in fluorescence intensity is most likely due to Ca^2+^ entering through the mechanotransduction channel (Zhang et al., 2018).

**Figure 6.**
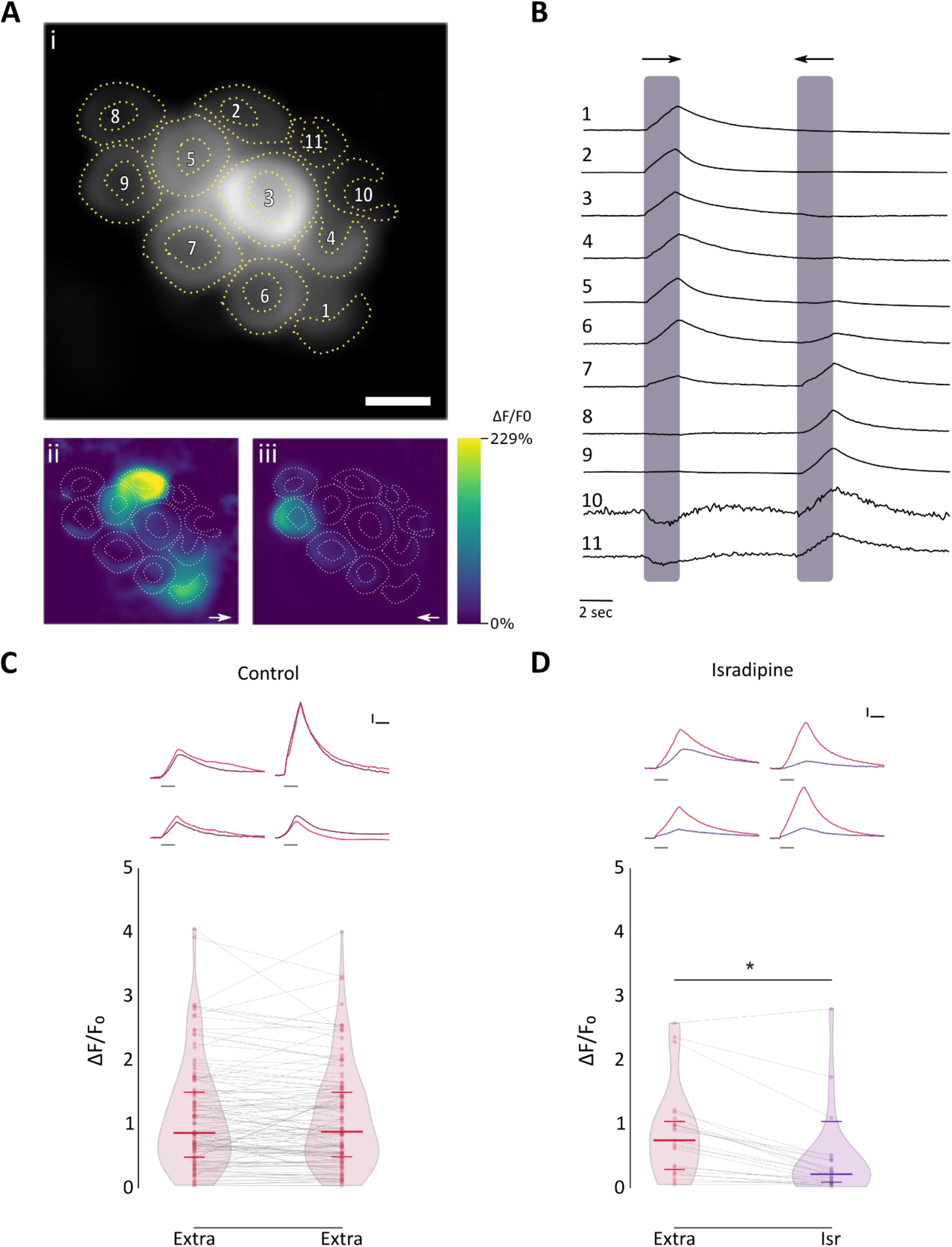
Effect of isradipine on mechanosensitive Ca^2+^ signals. **A**, Representative functional Ca^2+^ images of a double transgenic neuromast expressing GcAMP7a in HC. **i** Pre-stimulus baseline grayscale image (ROIs are drawn around each visible hair cell), **ii** and **iii** Spatial patterns of GCaMP7a Ca^2+^ signals, during a 2 sec mechanical stimulus in either the anterior-posterior (→) or in the posterior-anterior direction (←), are color coded according to the ΔF/F0 heat map. **B**, Representative temporal curves of mechanosensitive Ca^2+^ responses (ΔF/F0) of HC numbered in **A**, normalized to the peak intensity for each cell. Shaded areas indicate the time when the neuromast was mechanically stimulated. **C**, *Top*, Representative temporal curves of mechanosensitive Ca^2+^ responses of 4 HC, over two trials with same stimulation after 1 minute (1° stimulus (light red), 2° stimulus (dark red)). *Bottom*, Peak ΔF/F0 for single HC (n = 113, each in its preferred orientation) over two trials with the same stimulation 1 minute apart. **D**, *Top*, Representative temporal curves of mechanosensitive Ca^2+^ responses of 4 HC, before (red) and after (purple) pre-incubation with 10 μM isradipine. *Bottom*, Pre-incubation with 10 μM isradipine drastically reduced peak ΔF/F0 (n = 23, W= −258, *p = 8.726e-05, MPRBC= 0.935; Wilcoxon matched-pairs signed rank test). Extra: Extracellular imaging solution, Isr: Isradipine. Scale bar in **A**: 5 μm. Horizontal scale bar in **C** and **D:**1.5 sec, Vertical scale bar in **C** and **D**: 25% ΔF/F0. Duration of the stimulus in **C** and **D***Top* is indicated by gray lines below each trace.

Stimulation of cholinergic efferents in the LL of *Xenopus*, burbot *Lota lota*, and dogfish *Scyliorhynus*, inhibits spontaneous and evoked activity of afferents by generating inhibitory postsynaptic potentials in HC (Russell Ij, 1971; Roberts and Russell, 1972; Flock and Russell, 1976). In zebrafish, activation of cholinergic efferents suppresses glutamate release from HC (Pichler and Lagnado, 2020) and inhibits afferent activity (Lunsford et al., 2019). This most likely results from ACh-evoked hyperpolarization of HC and reduced Ca^2+^ influx through voltage-activated Ca^2+^ channels.

In order to analyze the effect of nAChR activation on HC, we evaluated the change in fluorescence intensity on mechanically-stimulated HC pretreated with ACh. If the activation of nAChR leads to HC hyperpolarization, then exogenous application of ACh should result in a decreased change in fluorescence intensity due to a reduced activation of Cav1.3 channels and thus lower Ca^2+^ influx. As expected, exogenous application of 1 mM ACh on mechanically-stimulated HC elicited a significant decrease in evoked Ca^2+^ influx with respect to control (Figure 7A and B, Ctrl: Mdn ΔF/F0= 0.818 IQR: 0.341-1.479 Vs. ACh: Mdn ΔF/F0= 0.573 IQR: 0.315-1.101, n = 114, W= −3493, p = 7.89e-07, MPRBC = 0.532, Wilcoxon matched-pairs signed rank test). This effect was reversed by superfusing the preparation with extracellular imaging solution (Figure7C, n= 37, Friedman test, Q= 18.54, p=9.418e-05. Dunn’s multiple comparisons test, Extra Vs. ACh: p=0.000705, Extra Vs. Wash: p=0.608054). ACh application *per se* did not evoke changes in basal fluorescence intensity (Figure 7D, naïve: 655.9 ± 81.1 Arbitrary Units (A.U.) vs. ACh: 652.2 ± 78.55 A.U, n = 45 cells, t= 0.7816, df= 44, p = 0.4386, two-tailed paired t-test). Furthermore, ACh mediated effect was observed in HC irrespective of polarity. Figure 7E shows that exogenous application of ACh elicited a significant decrease in mechanically-evoked Ca^2+^ influx, both in HC selectively activated by an anterior to posterior (Ant-Post) or by a posterior to anterior (Post-Ant) stimulus (Ant-Post, Extra Mdn ΔF/F0= 0.783 - ACh Mdn ΔF/F0= 0.559, n= 62, W= −1227, p= 1.698e-05, MPRBC= 0.628; Post-Ant, Extra Mdn ΔF/F0= 0.831 - ACh Mdn ΔF/F0=0.653, n= 52, W= −582, p=0.008, MPRBC= 0.422; Wilcoxon matched-pairs signed rank test). In addition, there was no significant difference in the magnitude of ACh mediated-effect between HC with different polarity selectivity (Figure 7F, (Ant-Post Mdn rel. ΔF/F0 diff = −0.2003 IQR: −0.424-0.051 Vs. Post-Ant Mdn rel. ΔF/F0 diff = −0.241 IQR: −0.497-0.118, U= 1560, p = 0.7687; Mann-Whitney test). Noteworthy, ACh-mediated effect on evoked Ca^2+^ signals was heterogeneous. To quantify the degree of inhibition elicited by ACh we used an *ad-hoc* metric, the Inhibition Index (*II*), that was calculated for each HC in Figure 7A. If ΔF/F0_extra_ is the change in fluorescence intensity on mechanically-stimulated HC under control conditions and ΔF/F0_ACh_ is the change in fluorescence intensity on mechanically-stimulated HC pretreated with ACh, then *II* was calculated as:

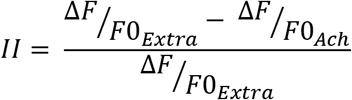

**Figure 7.**
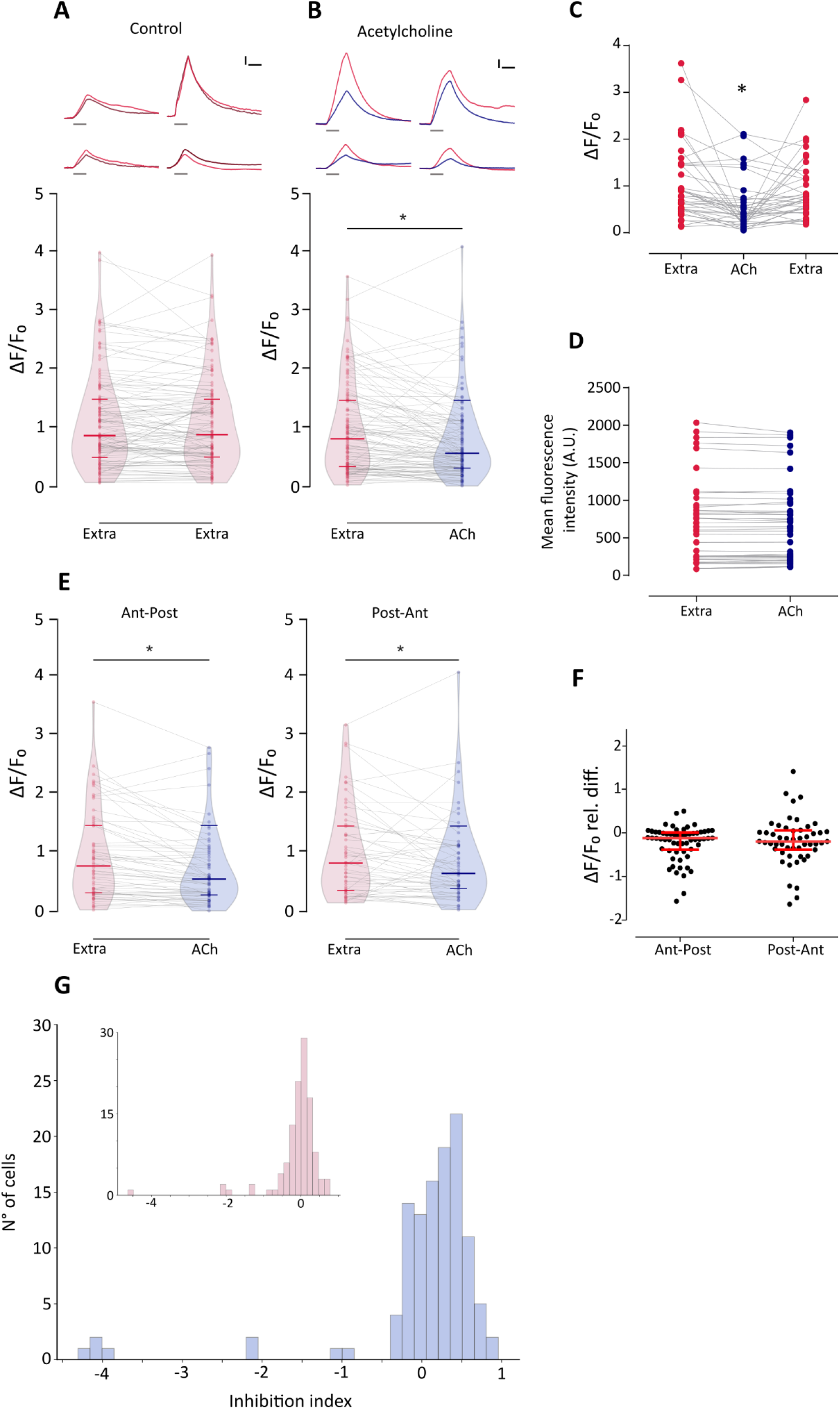
ACh inhibits mechanically-evoked Ca^2+^ signals and this inhibition is heterogeneous and independent of HC polarity **A**, *Top*, Representative temporal curves of mechanosensitive Ca^2+^ responses of 4 HC, over two trials with the same stimulation 1 minute apart (1° stimulus (light red), 2° stimulus (dark red)). *Bottom*, Peak ΔF/F0 for single HC (n = 113) over two trials with the same stimulation after 1 minute. **B**, *Top*, Representative temporal curves of mechanosensitive Ca^2+^ responses of 4 HC, before (red) and after (blue) the application of 1 mM ACh. *Bottom*, ACh application reduces mechanosensitive Ca^2+^ responses (n = 114, W= −3493, *p = 7.89e-07, MPRBC = 0.532, Wilcoxon matched-pairs signed rank test). **C**, ACh-mediated reduction in mechanically-evoked Ca^2+^ signals is reversed after 1 minute wash with extracellular imaging solution. (n= 37, Friedman test, F= 18.54, p=9.418e-05, Dunn’s multiple comparisons test, Extra Vs. ACh: p=0.000705, Extra Vs. Wash: p=0.608054). **D**, Basal Ca^2+^ levels show no significant differences before and after the application of 1 mM ACh (n = 45 cells, t= 0.7816, df= 44, p = 0.4386, two-tailed paired t-test). **E**, ACh reduces mechanosensitive Ca^2+^ responses in HC of opposing polarity (Ant-Post, n= 62, W=−1227, *p= 1.698e-05, MPRBC= 0.628; Post-Ant, n= 52, W= −582, *p=0.008, MPRBC= 0.422; Wilcoxon matched-pairs signed rank test). **F**, HC of opposing polarity exhibit no significant differences between their ACh-mediated relative change in peak ΔF/F0. (U= 1560, p = 0.7687; Mann-Whitney test). **G**, Distribution of Inhibition Index (*II*, calculated as (ΔF/F0extra −ΔF/F0ACh) / ΔF/F0extra) for ACh-treated HC. *Inset*, Distribution of Change Index (*CI*, calculated as (ΔF/F0stim1 −ΔF/F0stim2) / ΔF/F0stim1) for two successive mechanical stimuli under control conditions. The distribution of *CI* is centered around 0. A reduced number of cells (<10%) exhibit large negative *CI* values that occur when fluorescence signal is greater during the 2° stimulus, suggesting these might be outliers. Lines indicate the median and IQR. Extra: Extracellular imaging solution, ACh: Acetylcholine, Ant - Post: Anterior - Posterior deflection preference. Post - Ant: Posterior - anterior deflection preference. Horizontal scale bar in **A** and **B**: 1.5 sec, Vertical scale bar in **A** and **B**: 25% ΔF/F0. Duration of the stimulus in **A** and **B***Top* is indicated by gray lines below each trace.

Thus, *II*=0 indicates no inhibition, *II*=1 indicates full inhibition and 0 >*II*>1 indicates partial inhibition. Figure 7G shows the distribution of *II* for ACh-treated HC. As expected, in the majority of cases *II* values were > 0. However, a subpopulation of cells exhibited *II* values close to 0, denoting no ACh-mediated inhibition. The absence of ACh-treated HC with *II*=1 is a consequence of our selection criteria, that is selecting cells with measurable mechanically-evoked Ca^2+^ signals in both control conditions and during ACh treatment. Consequently, the inhibitory effect of ACh might be underestimated.

To assess the identity of the nAChR mediating synaptic transmission at the LL efferent synapse, we tested the effect of α-Btx, a potent inhibitor of recombinant zebrafish α9 and α9α10 nAChRs, on ACh-mediated inhibition of evoked Ca^2+^ signals. Figure 8B shows that ACh modulation was blocked when this agonist was co-applied with 10 μM α-Btx (α-Btx: Mdn ΔF/F0= 0.534 IQR: 0.3308-1.325 Vs. ACh-α-Btx: Mdn ΔF/F0= 0.4015 IQR: 0.203-0.816, n = 25, W= −87, p= 0.2541, MPRBC = 0.268, Wilcoxon matched-pairs signed rank test), supporting the hypothesis that a functional α9* nAChR is present at the zebrafish LL efferent synapse.

**Figure 8.**
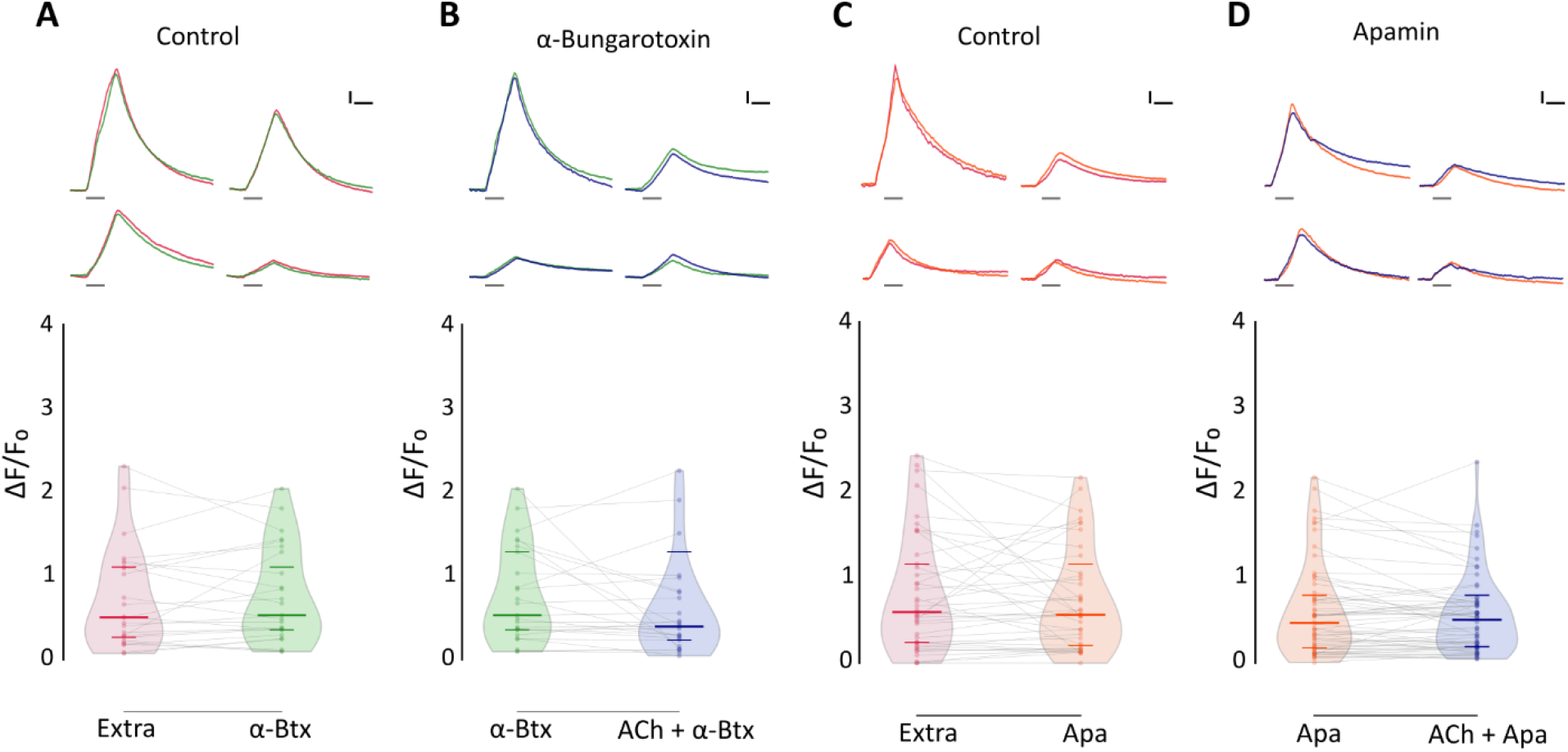
ACh-mediated inhibition of evoked Ca^2+^ signals is blocked by α-Btx and apamin. **A**, *Top*, Representative temporal curves of mechanosensitive Ca^2+^ responses of 4 HC, before (red) and after the application of 10 μM α-Btx (green). *Bottom*, Mechanosensitive Ca^2+^ signals show no significant difference before and after 10 μM α-Btx treatment (Extra: Mdn ΔF/F0= 0.509 IQR: 0.252-1.134 Vs. α-Btx: Mdn ΔF/F0= 0.534 IQR: 0.331-1.325, n = 25, W= −45, p= 0.5449, MPRBC = 0.138). **B**, *Top*, Representative temporal curves of mechanosensitive Ca^2+^ responses of 4 HC, after the application of 10 μM α-Btx (green) and after the co-application of 1 mM ACh and 10 μM α-Btx (blue). *Bottom*, When co-applied with 10 μM α-Btx, ACh-mediated inhibition is blocked (n = 25, W= −87, p= 0.2521, MPRBC = 0.268). **C**, *Top*, Representative temporal curves of mechanosensitive Ca^2+^ responses of 4 hair cells, before (red) and after the application of 10 μM apamin (orange). *Bottom*, Mechanosensitive Ca^2+^ signals show no significant difference before and after 10 μM apamin treatment (Extra: Mdn ΔF/F0= 0.599 IQR: 0.243-1.216 Vs. Apa: Mdn ΔF/F0= 0.567 IQR: 0.191-1.040, n = 41, W= −91, p= 0.5554, MPRBC = 0.106). **D**, *Top*, Representative temporal curves of mechanosensitive Ca^2+^ responses of 4 HC, after the application of 10 μM apamin (orange) and after the co-application of 1 mM ACh and 10 μM apamin (blue). *Bottom*, ACh-mediated inhibition is blocked by 10 μM apamin (n = 60, W= −322, p= 0.2359, MPRBC = 0.1759). A Wilcoxon matched-pairs signed rank test was used in all cases. Extra: Extracellular imaging solution, ACh: Acetylcholine, α-Btx: α-bungarotoxin, Apa: Apamin. Horizontal scale bar in **A**, **B**, **C** and **D**: 1.5 sec, vertical scale bar in **A**, **B**, **C** and **D**: 25% ΔF/F0. Duration of the stimulus is indicated by gray lines below each trace.

In mammals and birds, the inhibitory sign of the efferent synapse is due to the entry of Ca^2+^ through α9α10 receptors and the subsequent activation of a small-conductance SK2 Ca^2+^-dependent potassium channel (Hiel et al., 2000; Oliver et al., 2000; Gómez-Casati et al., 2005; Matthews et al., 2005; Elgoyhen and Katz, 2012). To evaluate the coupling of ACh responses to SK activation in zebrafish LL efferent synapse, we analyzed the effect of apamin, a known SK channel blocker, on ACh-mediated inhibition of evoked Ca^2+^ signals. Co-application of 1 mM ACh and 10 μM apamin abolished the inhibitory effect of ACh (Figure 8D, Apa: Mdn ΔF/F0= 0.470 IQR: 0.176-0.818 Vs. Apa-α-Btx: Mdn ΔF/F0= 0.508 IQR: 0.192-0.718, n = 60, W= −322, p=0.2359, MPRBC = 0.1759, Wilcoxon matched-pairs signed rank test), suggesting that the nAChR that serves the LL efferent synapse is functionally coupled to an SK channel.

## Discussion

Vertebrate HC systems are innervated by descending efferent fibers that modulate their response to external stimuli (Russell, 1971; Metcalfe et al., 1985; Guinan and Stankovic, 1996; Bricaud et al., 2001). In the LL, the excitation of efferent fibers inhibits afferent activity by generating inhibitory postsynaptic potentials in HC (Russell, 1971; Flock and Russell, 1973, 1976). In addition, excitatory efferent effects can be observed when cholinergic transmission is blocked (Flock and Russell, 1973). The latter is mediated by dopamine released in a paracrine fashion from efferent fibers located within the supporting cell layer and acting through D1b receptors (Toro et al., 2015). To date, the nature of the molecular players for ACh-mediated inhibitory effects have remained unknown. Here we provide evidence for a mechanism by which an α9* nAChR operates at the zebrafish LL efferent synapse. Our study suggests that upon ACh release, Ca^2+^ influx through these receptors activates nearby SK channels, leading to LL HC hyperpolarization (Figure 9).

**Figure 9.**
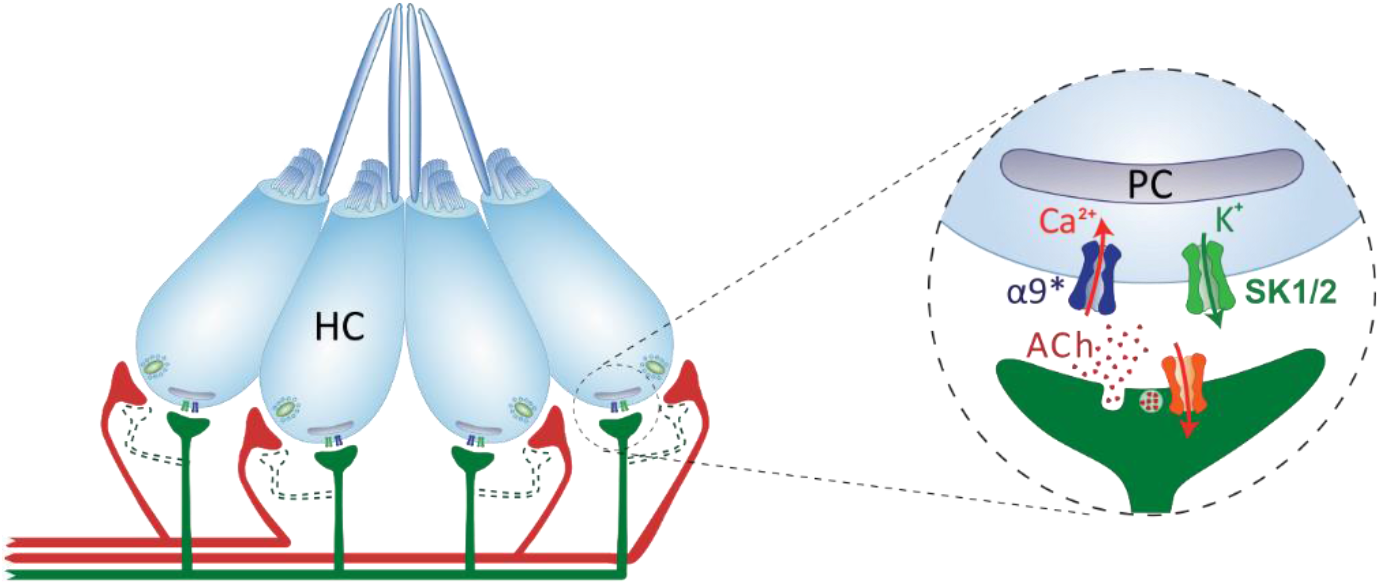
Schematics of the cholinergic LL efferent synapse. LL HC are innervated by afferent (red) and cholinergic efferent (green) fibers. Evidence for efferent cholinergic fibers contacting afferent neurons (dashed light green) is still missing. The net effect of LL efferent cholinergic activity is to hyperpolarize HC. This is mediated by the activation of an α9* nAChR with high Ca^2+^ permeability. Subsequent activation of Ca^2+^-dependent potassium SK channels drives HC hyperpolarization. Postsynaptic cisterns (PS) opposed to efferent terminals (Dow et al., 2018) have been proposed to participate in Ca^2+^ compartmentalization and/or Ca^2+^ induced Ca^2+^ release mechanisms.

Efferent innervation mediated by α9* nAChRs is a common feature to all vertebrate HC (Elgoyhen et al., 1994; Glowatzki and Fuchs, 2000; Hiel et al., 2000; Holt et al., 2003; Parks et al., 2017). In mammals, MOC efferent activity is mediated by α9α10 nAChRs (Elgoyhen et al., 2001; Lustig et al., 2001; Sgard et al., 2002; Gómez-Casati et al., 2005). In fact, α10 subunits are strictly required for efferent function, since in α10 knockout mice, α9 nAChRs expressed by OHC are unable to transduce efferent signals *in vivo* (Vetter et al., 2007). Surprisingly, our analysis of single-cell expression studies showed enriched expression of α9 but not α10 subunits in zebrafish HC. This is in accordance with Lush et al. (2019) who have shown that chrna9 but not chrna10 is differentially expressed in mature LL HC. Moreover, the present *in situ* hybridization data indicates expression of α9 of but not α10 mRNA in neuromast HC. In addition, our functional data proved that zebrafish α9 nAChRs expressed in *Xenopus* oocytes are functional and exhibit robust ACh-evoked currents which are not boosted in magnitude when co-expressed with α10. This is in stark contrast to that observed for rat α9 receptors which, although functional when heterologously expressed, exhibit very small ACh-evoked responses, which are non-reliable nor reproducible, and significantly boosted when co-expressed with α10 (Elgoyhen et al., 1994, 2001; Sgard et al., 2002). Moreover, the EC_50_ for ACh of zebrafish α9α10 nAChRs is 40-times higher than that of α9 homomeric receptors and near 500 **μ**M, a value that is too high compared to any other known α9* nAChR EC_50_ (Elgoyhen et al., 2001; Lipovsek et al., 2012; Marcovich et al., 2020). Taken together, our expression and functional data strongly suggest that the nAChR at the LL efferent synapse is an α9 homomeric receptor.

Striking features of the zebrafish α9 nAChR are its high desensitization rate and lack of modulation by external Ca^2+^. This differs from that reported for rat (Elgoyhen et al., 1994; Katz et al., 2000) and chicken (Lipovsek et al., 2012) α9 homomeric receptors, which exhibit low desensitization kinetics, and in the case of rat receptors (not reported for chicken), are blocked by extracellular Ca^2+^. These results support the observation that within the nAChR family, α9 and α10 subunits are the ones that exhibit the highest degree of coding sequence divergence, mirrored by a great variability of functional properties across species (Franchini and Elgoyhen, 2006; Lipovsek et al., 2012; Marcovich et al., 2020). Thus, as reported by Marcovich et al.(2020), our results further indicate that differences in efferent modulation of HC activity across species could have shaped the functional properties of α9* receptors and vice versa.

The high desensitization kinetics of zebrafish α9* receptors leads to a self-limiting Ca^2+^ entry through this highly Ca^2+^ permeable nAChR. This is key to prevent crosstalk between efferent and afferent systems, which co-exist in LL HC, and could lead to Ca^2+^ spillover from efferent-mediated Ca^2+^ entry to Ca^2+^-triggered glutamate release and activation of afferent fibers, as reported in developing inner HC (Moglie et al., 2018). Furthermore, postsynaptic cisterns opposed to LL efferent terminals (Dow et al., 2018) could also provide efficient compartmentalization of cholinergic Ca^2+^ signals to prevent efferent-to-afferent cross-talk. In rat and chicken, the desensitization capability of α9α10 receptors is provided by the α10 subunit (Elgoyhen et al., 2001; Lipovsek et al., 2012). Since in zebrafish LL the efferent response most likely relies on α9 nAChRs, one could propose that non-synonymous substitutions in the coding sequence of this subunit might lead to a receptor highly fitted to convey self-limiting Ca^2+^ influx into HC, and this should be further tested.

The increase in intracellular Ca^2+^ upon deflection of the cilia, results from Ca^2+^ influx through mechanosensitive ion channels (Corey and Hudspeth, 1979; Fettiplace, 2009; Zhang et al., 2018) and the subsequent activation of voltage-gated Ca^2+^ channels due to HC depolarization (Moser and Beutner, 2000; Sheets et al., 2017; Zhang et al., 2018). The fact that the application of ACh resulted in a reduction of Ca^2+^ signals, most likely indicates a reduced depolarization. Thus one could propose that ACh inhibits Ca^2+^ influx due to a net hyperpolarization of LL HC. The finding that ACh-mediated effect can be blocked by apamin, supports the generally held hypothesis that Ca^2+^ entering through the efferent nAChR activates nearby SK channels leading to HC hyperpolarization (Doi and Ohmori, 1993; Blanchet et al., 1996; Nenov et al., 1996; Yuhas and Fuchs, 1999; Glowatzki and Fuchs, 2000; Oliver et al., 2000; Holt et al., 2003, 2003; Katz et al., 2004; Dawkins et al., 2005; Gómez-Casati et al., 2005; Parks et al., 2017). The SK2 nature of the K^+^ channel functionally coupled to α9* nAChRs has been established in birds (Matthews et al., 2005) and mammals (Dulon et al., 1998; Oliver et al., 2000). However our cross-study analysis revealed that both kcnn2 (SK2) and kcnn1b (SK1b) transcripts are enriched in zebrafish HC. This is consistent with Cabo et al. (2013) showing SK1 expression in zebrafish LL HC, but not in ear sensory epithelia. Interestingly, SK1 and SK2 are generally co-expressed in the brain of the electric fish *Apteronotus leptorhynchus* (Ellis et al., 2008) and mammals (Stocker and Pedarzani, 2000). Moreover, rat SK1 forms functional heteromeric channels with SK2 (Benton et al., 2003; Autuori et al., 2019). The fact that apamin blocked ACh-mediated effects suggests that SK2 channels and not SK1 play a key role in zebrafish LL HC hyperpolarization, since SK2 channels are the most apamin-sensitive (Köhler et al., 1996; Shah and Haylett, 2000; Strøbaek et al., 2000; Stocker, 2004).

Zebrafish neuromasts contain two populations of HC which are activated by deflections in either the anterior or posterior direction (Flock and Wersall, 1962; Ghysen and Dambly-Chaudiere, 2007). However, only one efferent fiber contacts all HC of a single neuromast (Faucherre et al., 2009; Dow et al., 2018). Moreover, during fictive locomotion presynaptic activity across all efferent synapses within a neuromast are synchronously activated (Pichler and Lagnado, 2020). Similarly, in our experiments ACh-mediated effects were observed in HC irrespective of their polarity and the median magnitude of inhibition in both cases showed no significant difference. However, it has been reported that efferent modulation is biased toward HC activated during forward motion (Pichler and Lagnado, 2020). This discrepancy might relay on the fact that our experiments were performed by the perfusion of ACh and not by stimulation of efferent terminals, which would better resemble experiments reported by Pichler and Lagnado (2020). Differences in the efficiency of presynaptic ACh release at efferent terminals and/or in the number of efferent terminals per HC of different polarities might account for this biased efferent modulation. Alternatively, physiological heterogeneity of LL HC could contribute to differences in the efficiency with which depolarization triggers glutamate release.

HC within the same neuromast exhibit functional heterogeneity, since stimuli able to open mechanosensitive channels are insufficient to evoke vesicle fusion in the majority of HC (Zhang et al., 2018). It has been shown that synaptically active HC exhibit lower intracellular K^+^ levels than silent HC. Moreover, physiological heterogeneity has also been shown for LL afferent response to efferent activity (Lunsford et al., 2019). In tune with these findings, we show that ACh-mediated effect on evoked Ca^2+^ signals is heterogeneous, adding a new level of complexity underlying LL HC function *in vivo*. Differences in the density of α9* nAChRs mediating Ca^2+^ influx and/or SK channels causing hyperpolarization could explain this phenomenon.

The LL system controls many behaviors such as obstacle and predator avoidance, schooling (Partridge and Pitcher, 1980; Mekdara et al., 2018), prey capture (McHenry et al., 2009; Stewart et al., 2013) and rheotaxis (Bleckmann and Zelick, 2009; Olszewski et al., 2012; Suli et al., 2012; Oteiza et al., 2017). However, the role of the efferent system on the performance of these behaviors is still unknown. Deciphering the molecular players at the zebrafish cholinergic LL efferent synapse is a step forward that enables the generation of molecular tools to selectively manipulate its activity and evaluate its role on sensory processing and associated behaviors in their native context.

## Contributions

ACF, MM, PVP and ABE designed research, ACF, MM, TC, LS and CW performed research, IM, SG, SD and JCG contributed unpublished reagents/analytic tools, ACF and PVP analyzed data, ACF, PVP and ABE wrote the paper.

## Acknowledgments

This research was supported by Agencia Nacional de Promoción Científica y Técnica (Argentina) (PICT 2016-2537 to ABE and PICT 2013-1117 to PVP), Consejo Nacional de Investigaciones Científicas y Técnicas (Argentina) (PIP 2014-301 to PVP), Human Frontiers in Science Program (RGP0033/2014 to Hernán Lopez-Schier, Florian Engert and ABE) and Scientific Grand Prize from the Fondation Pour L’Audition to ABE. The authors would like to thank Hernán Lopez Schier and Florian Engert for academic discussions.

---

**Figure 1-1.**
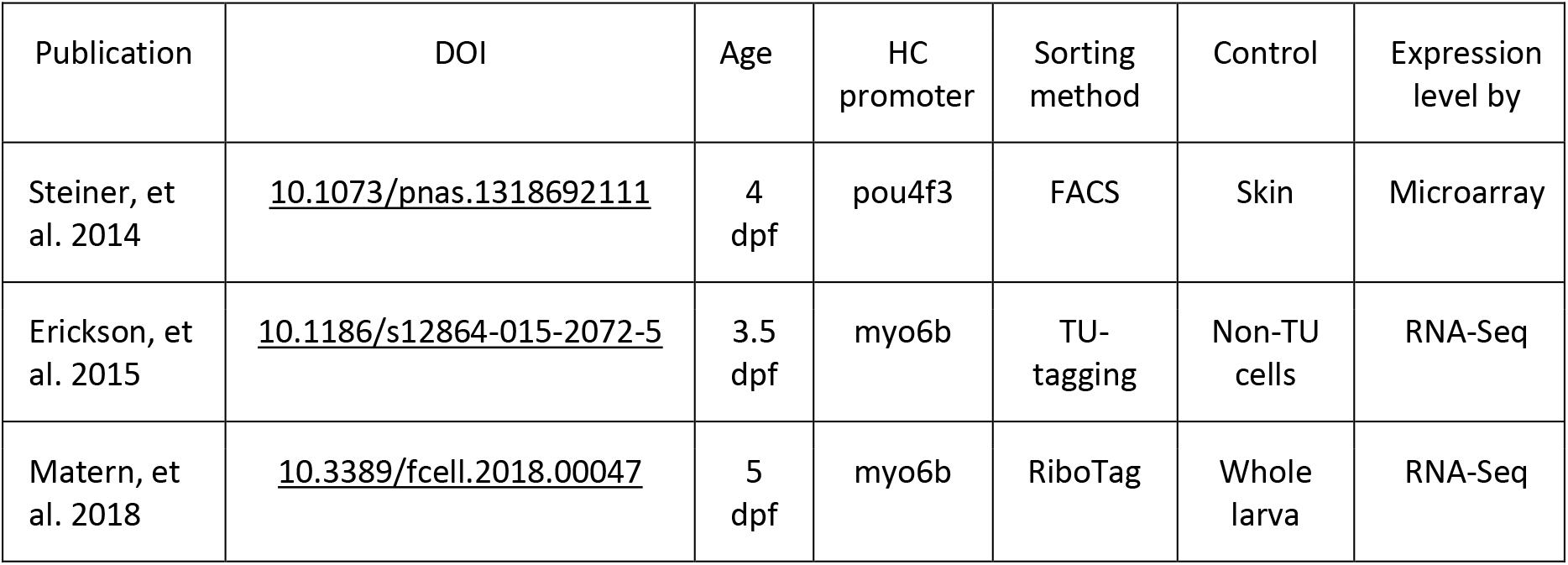
Meta-data of the studies used to analyze gene enrichment in zebrafish HC

**Figure 1-2.**
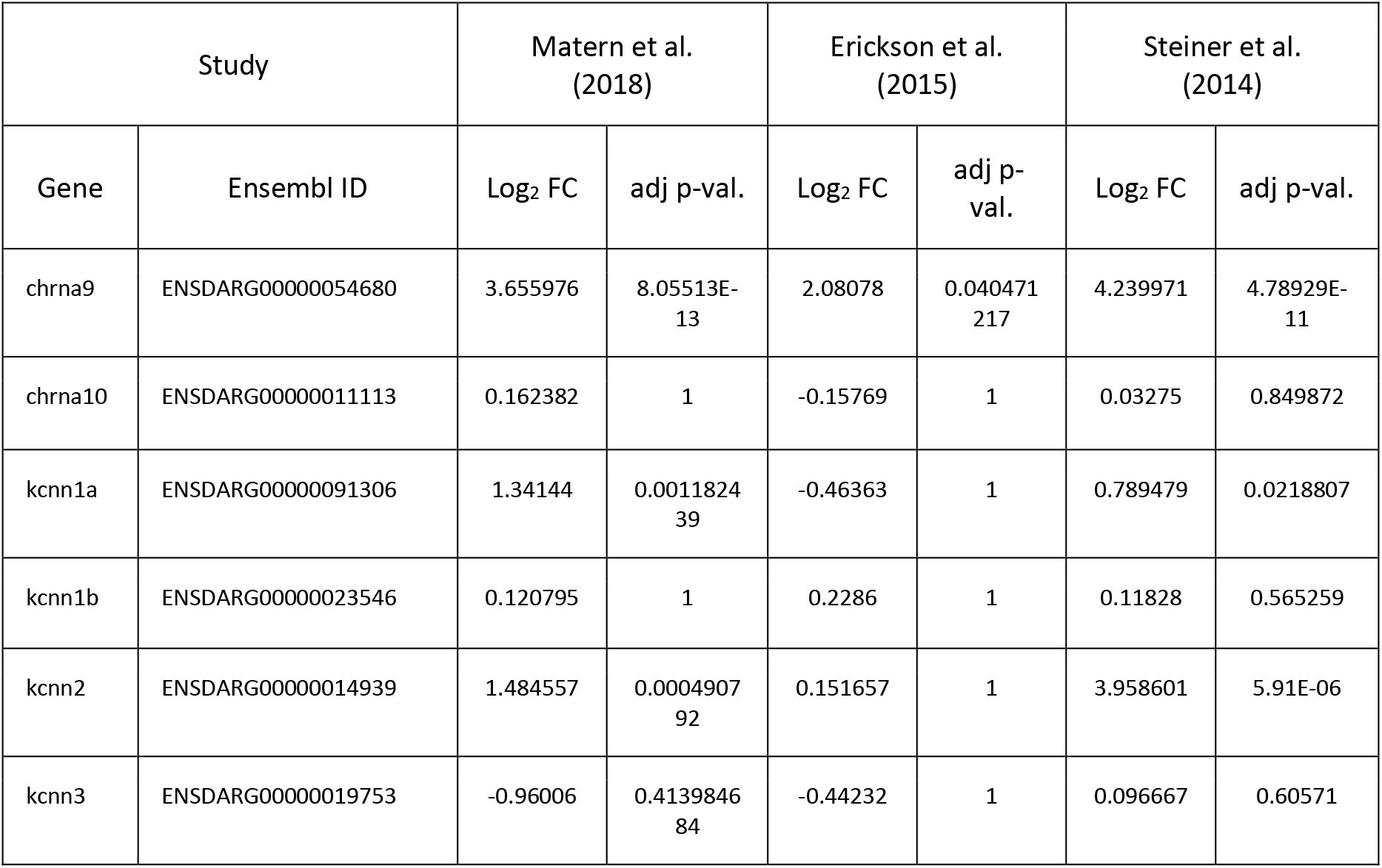
Log_2_ Fold Change and adjusted p-values for the genes of interest across the different studies analyzed.

